# Taking the short- or long-chain route: conversion efficiency of alpha linolenic acid to long-chain omega-3 fatty acids in aerial insectivore chicks

**DOI:** 10.1101/155499

**Authors:** Cornelia W. Twining, Peter Lawrence, David W. Winkler, Alexander S. Flecker, J. Thomas Brenna

## Abstract

Food availability and quality are both critical for growing young animals. In nature, swallows (*Tachycineta bicolor*) and other aerial insectivores feed on both aquatic insects, which are rich in omega-3 long-chain polyunsaturated fatty acid (LCPUFA) and terrestrial insects, which contain considerably less LCPUFA. Carnivorous mammals and fishes must obtain LCPUFA from diet, as they have lost the capacity to convert the precursor omega-3 ALA into LCPUFA. Thus, the relative value of aquatic versus terrestrial insects depends not only on the fatty acid composition of the prey, but also upon the capacity of consumers to convert ALA into LCPUFA. We used a combination of stable-isotope-labeled fatty acid tracers to ask if, and how efficiently, Tree Swallows can deposit newly synthesized LCPUFA into tissue. Our data show for the first time that Tree Swallows can convert ALA into LCPUFA deposited in liver and skeletal muscle. However, high Tree Swallow demand for LCPUFA combined with low ALA availability in natural terrestrial foods may strain their modest conversion ability. This suggests that while Tree Swallows can synthesize LCPUFA de novo, LCPUFA are ecologically essential nutrients in natural systems. Our findings thus provide mechanistic support for our previous findings and the importance of LCPUFA-rich aquatic insects for Tree Swallows and most likely other aerial insectivores with similar niches.

**Summary Statement:** A stable-isotope-labeled tracer reveals the mechanism for omega-3 long-chain polyunsaturated fatty acid (LCPUFA) limitation in a wild avian insectivore, showing that LCPUFA are an ecologically essential nutrient.

## Introduction

Although dietary resources are crucial for animals throughout their life cycle, energy and nutrients are particularly critical during early life, especially for animals, like birds, that undergo rapid determinate growth. For wild birds with young that reach mature size rapidly, timing breeding phenology with food availability is essential for survival (Lyon et al. 2008). Climate change is already creating mismatches between the phenology of insect prey and the breeding timing of insectivores like Pied Flycatchers (*Ficedula hypoleuca*; Both et al. 2006) and Great Tits (*Parus major*; Nussey et al. 2005). However, even if food resources are available during the breeding season, food availability in terms of energy alone may not be sufficient for successful breeding success if available foods lack key nutrients. Recent studies suggest that there is also the potential for mismatches between the nutritional composition of available food resources and the complex nutritional needs of growing chicks (Twining et al. 2016a; Twining et al. submitted).

Food quality can be defined in many ways, including caloric density, nutrient composition, and digestibility. Here, we focus on food quality in terms of the availability of long-chain polyunsaturated omega-3 fatty acids (LCPUFAs), which we previously found to be more important for insectivore chick growth performance than was food availability (Twining et al. 2016a). LCPUFAs, in particular the fatty acids DHA (docosahexaenoic acid, 22:6n-3) and EPA (eicosapentaenoic acid, 20:5n-3), are important organic compounds for most animals: they affect a range of important physiological processes from immune function to vision and brain development (Twining et al. 2016b). However, some species, especially carnivores, are not able to synthesize LCPUFA. For example, cats, which are strict carnivores, and carnivorous fishes cannot synthesize any LCPUFA and must obtain them from their diets (Twining et al. 2016b). Omnivorous animals, from humans to many species of birds, can obtain LCPUFA through two pathways: 1) either directly by consuming food containing EPA and, DHA), or 2) indirectly by consuming their molecular precursor, the short chain omega-3 PUFA, alpha linolenic acid (18:3n-3, ALA), and then converting ALA into LCPUFA through the biochemical processes of elongation and desaturation (Brenna and Carlson 2014).

A major dichotomy in LCPUFA availability exists between aquatic and terrestrial ecosystems: LCPUFA are abundant at the base of aquatic food webs, and aquatic insects incorporate LCPUFA from aquatic primary producers into tissues (Twining et al. 2016b; Torres-Ruiz et al. 2007). In contrast, terrestrial plants contain little to no LCPUFA, though they contain their molecular precursor ALA (Hixson et al. 2015; Twining et al. 2016b), which is either incorporated into tissue or to a minor degree converted to LCPUFA by terrestrial insects (Blomquist et al. 1991). As a consequence, aquatic insects are much richer in LCPUFA than are terrestrial insects. In the wild, avian insectivores in riparian areas, such as Tree Swallows (*Tachycineta bicolor*), which are considered a model insectivore species, consume a mix of terrestrial and aquatic insect prey items (McCarty and Winkler 1999a; Winkler et al. 2013). Our recent studies show that differences in the fatty acid composition between aquatic and terrestrial insects can have strong consequences on Tree Swallow chick performance (Twining et al. 2016a) and breeding success (Twining et al. submitted).

Ultimately, the nutritional value of aquatic versus terrestrial insects depends not only on the fatty acid composition of these insects, but also upon the capacity of Tree Swallows and other insectivores to convert ALA into LCPUFA. While strict carnivores, such as cats, have lost the ability to elongate and desaturate ALA into LCPUFA and must obtain LCPUFA directly from diet, domestic chickens (*Gallus domesticus*), which are omnivorous, but consume mainly plant-based foods in captivity, appear to be relatively efficient at ALA to LCPUFA conversion (Twining et al. 2016b), with the capacity to survive and reproduce on LCPUFA-free diets. The majority of past studies on avian fatty acid requirements have focused on domesticated omnivorous taxa like chickens (e.g., Cherian and Sim 1991; Lin et al. 1991; Newman et al. 2002; Cherian et al. 2009). These studies found that domestic adult chickens and their chicks are capable of converting ALA to LCPUFA and that tissue LCPUFA increase with increased dietary ALA. Increasing ALA in maternal diet through foods, such as flax seeds, also increases LCPUFA content in eggs, embryos, and newly hatched chicks (Cherian and Sim 1991).

Studies on wild birds have found that both nutritional composition and foraging ecology affect avian tissue fatty acid composition and avian nutritional needs (McWilliams et al. 2002; Pierce et al. 2005; Maillet and Weber 2007; McCue et al. 2009). Most studies on wild birds (but see McWilliams et al. 2002; Pierce et al. 2005) have either looked at seed and fruit-eating passerines (e.g., McCue et al. 2009), which consume a diet much lower in LCPUFA, or fish- and shellfish-eating shorebirds, which consume a much higher LCPUFA diet than riparian aerial insectivores (e.g., McWilliams et al. 2004). Furthermore, past studies on avian fatty acid requirements have made inferences about conversion efficiency indirectly based on the effects of dietary fatty acid content on survival, performance, and tissue LCPUFA content, but they have not established whether ALA to LCPUFA conversion is metabolically possible, as have studies on humans and other mammals (Brenna et al. 2009).

Apart from obligate carnivores, animals generally retain the metabolic capacity to endogenously biosynthesize LCPUFA from precursors based in part on the LCPUFA content of their ancestral diets. Thus, terrestrial herbivores must endogenously synthesize effectively all of their LCPUFA while carnivores must obtain all the LCPUFA in their diets (Castro et al. 2012; Brenna and Carlson 2014; Twining et al. 2016a). On this basis, Tree Swallows and other riparian aerial insectivores should have limited capacity to convert ALA into the LCPUFA because they evolved with access to aquatic insects, obviating the need to maintain efficiency in this metabolic pathway. We previously found that LCPUFA have strong effects on Tree Swallow growth: chicks on high LCPUFA diets, which approximated diets dominated by aquatic insects, grew faster, were in better condition, had increased immunocompetence and decreased metabolic rates compared to chicks on low LCPUFA feeds approximating diets dominated by terrestrial insects (Twining et al. 2016a). We found similar patterns between dietary LCPUFA and chick performance in Eastern Phoebes (*Sayornis phoebe*; Twining et al. in prep). In a long-term field study, we also that found aquatic insects, which, unlike terrestrial insects, are rich in LCPUFA, are a strong driver of long-term Tree Swallow fledging success in nature (Twining et al. in prep). Together, these findings suggest that ALA to LCPUFA conversion, if present, is likely inefficient in Tree Swallows and other riparian aerial insectivore chicks.

To understand the physiological importance of LCPUFA for riparian aerial insectivores, we used rapidly growing Tree Swallow chicks as a model to ask: first, are birds that regularly consume high LCPUFA dietary resources able to convert ALA into LCPUFA or are dietary LCPUFA strictly essential nutrients? Second, if birds with access to high LCPUFA resources are able to convert ALA into LCPUFA, is their conversion and tissue deposition efficiency high enough to provide them with enough LCPUFA from dietary ALA in natural settings to avoid performance limitation or are dietary LCPUFA ecologically essential nutrients? We used enriched ^13^C stable isotope fatty acid tracers to trace the metabolic pathway of ALA through Tree Swallow chicks, adapting this methods from studies on humans and small mammals, to directly quantify ALA to LCPUFA conversion for the first time in wild birds.

## Methods

We examined ALA conversion capacity and efficiency in seven wild Tree Swallow chicks from two sites near Ithaca, New York (Site 1: 42.504434°N, 76.465949°W, Site 2: 42.515459°N, 76.335272°W). At both sites, we briefly removed chicks from the nest and fed them olive oil with or without dissolved δ13C-enriched ALA (Cambridge Isotope Laboratories, Cambridge, MA) via syringe. Six chicks were dosed with a δ13C-enriched ALA tracer, serving as treatment chicks, and one was not dosed with a δ13C-enriched ALA tracer and served as a natural abundance δ^13^C control. At the time of dosing, all chicks were approximately 7 days old and had a mean weight of 10.687 g (standard deviation = 1.204 g). We dissolved 5 mg of δ^13^C_ALA_ in 2.5 mL of olive oil creating a 10mg/mL solution of δ^13^C_ALA_ in olive oil. Each treatment chick received 0.25 mL of olive oil with dissolved δ^13^C_ALA_ followed by 0.25mL of olive oil without tracer using same syringe. The control chick received two 0.25 mL syringes of olive oil that did not come into contact with the tracer.

After dosing, we labeled chicks with non-toxic children’s nail varnish before returning them to their nests for parental care and feeding. All chicks were sacrificed at approximately 48 hrs post-dose per United States Fish and Wildlife Service migratory bird scientific collection permit #MB757670 and New York State Department of Environmental Conservation scientific collection permit #1477. All animal work was approved under Cornell Institutional Animal Care and Use Committee #2001-0051. After sacrificing chicks, we removed their livers and pectoral muscles for analyses.

Next, we performed compound-specific δ^13^C analysis and fatty acid composition analysis. Briefly, we extracted liver and pectoral muscle fatty acid methyl esters (FAMEs) using a modified one-step method (Garces and Mancha 1993). We quantified fatty acid composition using a BPX-70 (SGE Inc.) column and a HP5890 series II gas chromatograph-flame ionization detector (GC-FID). Chromatogram data were processed using PeakSimple. Response factors were calculated using the reference standard 462a (Nucheck prep). FAMEs were identified using a Varian Saturn 2000 ion trap with a Varian Star 3400 gas chromatography mass spectrometer run in chemical ionization mass spectrometry mode using Acteonitrile as reagent gas. We used gas chromatography combustion isotope ratio mass spectrometry (GCC-IRMS) to measure the δ^13^C signatures of ALA, EPA, and DHA (Goodman and Brenna 1992; Plourde et al 2014). Briefly, an Agilent 6890 GC was interfaced to a Thermo Scientific 253 isotope-ratio mass spectrometer via a custom-built combustion interface. Peaks were confirmed to be baseline separated, and calibrated against working standards with isotope ratios traceable to international standards calibrated to VPDB (Caimi et al. 1994; Zhang et al. 2009).

Conversion from ALA to LCPUFA requires a complex series of biochemical reactions of varying efficiency in each organ, including transport across and deposition into membranes. From measurement of the amount of labeled LCPUFA in tissue, we derive an apparent conversion efficiency (CE) reflecting all processes leading to the deposition of newly synthesized LCPUFA. This parameter, derived from experimental measurements, can also be understood as net deposition of labeled LCPUFA in tissue, rather than a mechanistic conversion rate that would be measured for a single enzymatic reaction. In experimental animals, this process is sensitive in the short-term to diet, with lower expression of genes involved in fatty acid conversion with LCPUFA feeding, as well as subject to competition due to the various fatty acids in the diet. We assessed ALA to EPA and DHA conversion efficiency by calculating conversion levels according to Sheaff et al. (1995). Atom Percent Excess (APE) as:
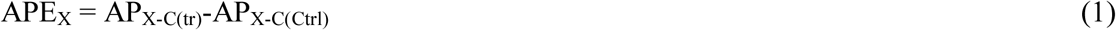

where AP are calculated directly from the measured δ^13^C_X_, X is a fatty acid (ALA, EPA, DPA, DHA), and tr and crtrl refer to chicked dosed with tracer (tr) or not dosed (Ctrl).

Total Label, or the concentration of tracer per unit weight of tissue, was calculated as:
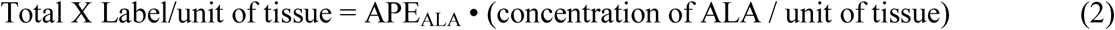

Finally, we calculated a tissue-specific apparent conversion efficiency (aCE) for fatty acids as:
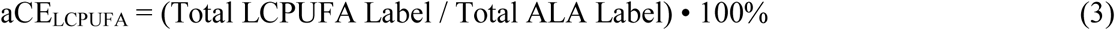

where LCPUFA refers to either DHA or EPA.

To understand how ALA-derived EPA and ALA-derived DHA accumulation varies between tissues where conversion activity occurs (i.e., liver) and those where it is expected to be lower (i.e., pectoral muscle), we divided liver CE by muscle CE for both EPA and DHA.

To understand the significance of our measured ALA conversion efficiency on performance, we applied our calculations of apparent CE to the mean ALA content of nestling bird feeds and wild aquatic and terrestrial insects to estimate potential EPA and DHA synthesis. We calculated potential EPA and DHA (LCPUFA) synthesis from ALA as:
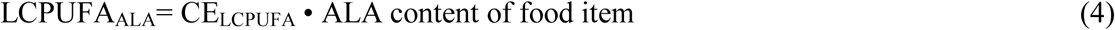

And potential total EPA and DHA (LCPUFA) as:
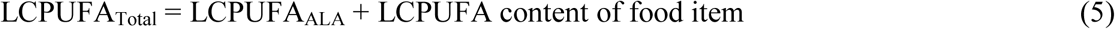

We then calculated the fraction of EPA and DHA (LCPUFA) from diet versus ALA conversion for each prey item as:
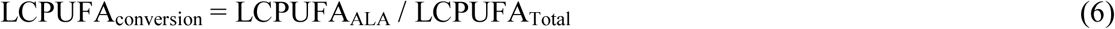

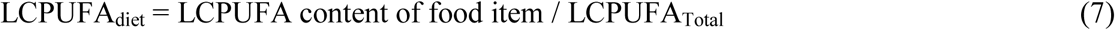

To understand how ALA-derived EPA and ALA-derived DHA accumulation varies between tissues where conversion activity occurs (i.e., liver) and those where it is minimal (i.e., pectoral muscle), we divided liver CE by muscle CE for both EPA and DHA.

To understand the potential impact of our measured ALA conversion efficiency on growth performance, we applied our calculations of conversion efficiency to the mean ALA content of nestling bird feeds and wild aquatic and terrestrial insects to estimate potential EPA and DHA synthesis. Insects were collected from eight sites around Ithaca, NY, using a combination of emergence traps, pan traps, and targeted hand-netting (Twining, unpublished). We determined insect fatty acid composition following the same methods described above for Tree Swallow tissue. We then calculated potential EPA and DHA (LCPUFA) synthesis from ALA as:
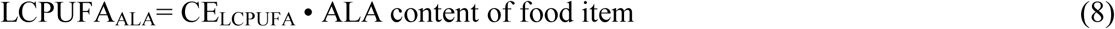

And potential total EPA and DHA (LCPUFA) as:
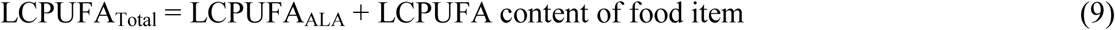

We then calculated the fraction of EPA and DHA (LCPUFA) from diet versus ALA conversion for each prey item as:
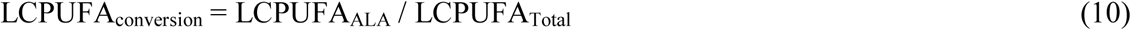

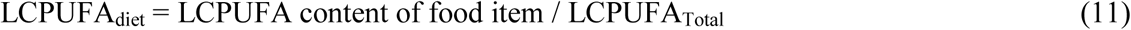

We calculated each of these measures for the following food items: 1) aquatic mayflies (Ephemeroptera Heptageniidae), 2) aquatic stoneflies (Plecoptera Perlidae), 3) aquatic dragonflies (Odonata Anisoptera), 4) terrestrial beetles (Coleoptera), 5) terrestrial flies (Diptera Brachycera), 6) terrestrial moths and butterflies (Lepidoptera), and 7) terrestrial bees (Hymenoptera Apidae).

We compared stable isotope values between our treatment chicks (n=6) and control chick (n=1) using one sample t-tests for both muscle and liver using control chick values as μ_0_. We used paired two-sample t-tests to compare conversion efficiency and tissue deposition between liver (n=7) and muscle (n=7) for treatment chicks. To compare raw ALA and EPA, potential ALA-derived EPA and DHA, and total potential EPA and DHA between aquatic and terrestrial insects, we also used two sample t-tests. We also used General Linear Models (GLM) to compare raw ALA and EPA, potential ALA-derived EPA and DHA, and total potential EPA and DHA between all insect groups using insect group as a factor. We performed post-hoc Tukey contrasts on all GLMs to determine which insect groups were significantly different from each other. All statistical analyses were performed in R version 3.3.3 (R core team).

## Results

We first asked if LCPUFA synthesis supports our previous results showing that LCPUFA are strictly essential nutrients for Tree Swallows. We found evidence that Tree Swallow chicks can derive LCPUFA from ALA: δ^13^C_EPA_ and δ^13^C_DHA_ values of liver and muscle from chicks fed δ^13^C-enriched ALA were significantly higher than controls (one sample t-tests: t-value = 4.62, df = 5, p-value < 0.01 for liver; t-value = 5.75, df = 5, p-value < 0.01; Table 1). Although all chicks fed δ^13^C-enriched ALA showed evidence of ALA to LCPUFA conversion and deposition of ALA-derived LCPUFA in tissues, we found substantial individual variation in conversion efficiency between individuals across both field sites, especially for DHA (Table 1). We found that ALA-derived EPA and ALA-derived DHA were significantly higher in liver, where most conversion activity is likely to occur, than in pectoral muscle, where LCPUFA are deposited (two sample t-test: t-value = 4.99, df = 5.42, p-value < 0.01 for EPA; t-value = 2.56, df = 5.23, p-value < 0.05 for DHA; Table 1).

**Table 1.** ALA to EPA and DHA conversion efficiency in Tree Swallow chick liver and pectoral muscle and ratio of liver to muscle conversion efficiency

| Tissue Type | Mean | Standard Deviation | Maximum | Minimum |
| --- | --- | --- | --- | --- |
| Liver EPA | 16.47% | 5.38% | 22.15% | 9.93% |
| Liver DHA | 18.06% | 13.76% | 34.05% | 0.23% |
| Muscle EPA | 5.32 % | 1.11% | 6.68% | 3.45% |
| Muscle DHA | 3.5% | 2.09% | 6.17% | 0.80% |
| Liver/Muscle EPA | 3.17 | 0.99 | 4.46 | 1.49 |
| Liver/Muscle DHA | 4.61 | 2.78 | 6.61 | 0.29 |

We next asked if LCPUFA are ecologically essential nutrients for Tree Swallows based on measured conversion efficiencies and tissue deposition levels combined with the fatty acid composition of potential insect prey (Table 2). Aquatic insects, especially mayflies and stoneflies, had much higher percentages of raw EPA than did terrestrial insects (two sample t-test for aquatic versus terrestrial EPA: t-value = 8.05, df = 13.96, p-value < 0.01; Table 3) while terrestrial insects, especially terrestrial moths, butterflies, and bees, had much higher percentages of ALA than did aquatic insects or terrestrial beetles and terrestrial flies (Figure 1a; two sample t-test for aquatic versus terrestrial ALA: t-value = -2.08, df = 42.81, p-value < 0.05; Table 3).

**Figure 1:**
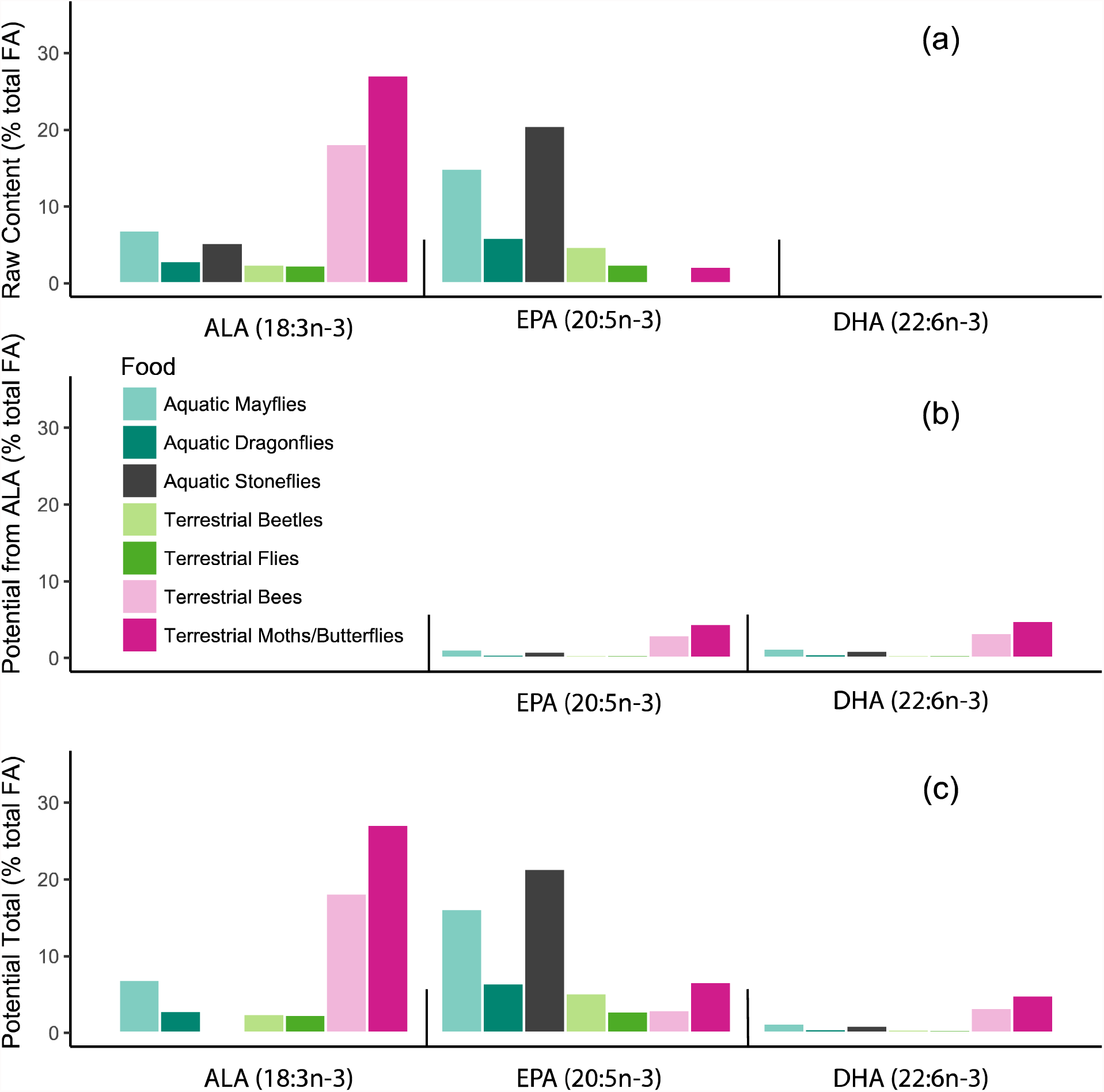
ALA, EPA, and DHA content of (a) food sources, (b) potential EPA and DHA from ALA to LCPUFA conversion, and (c) potential total ALA, EPA, and DHA content from food sources and ALC to LCPUFA conversion

**Table 2.**
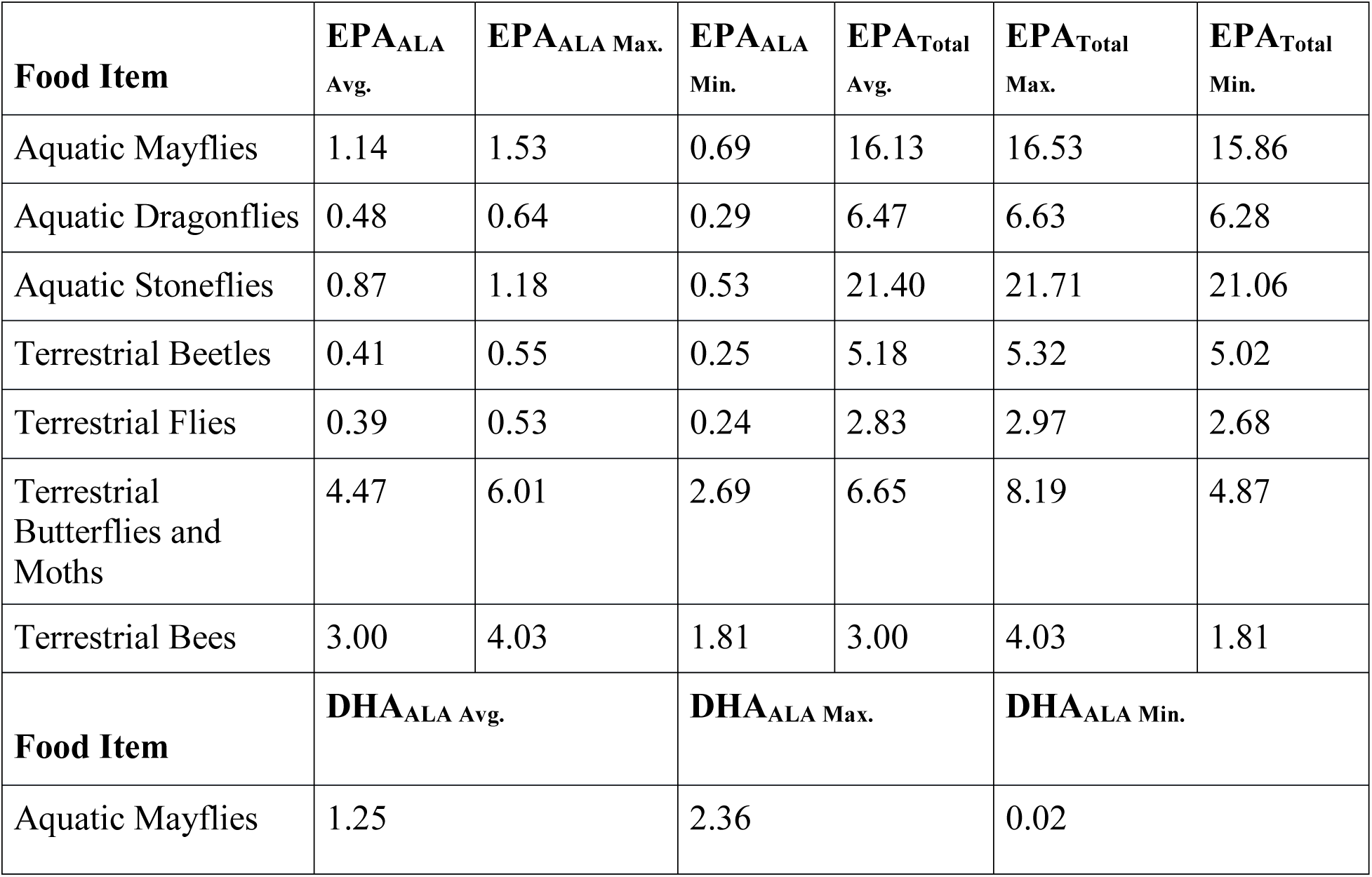
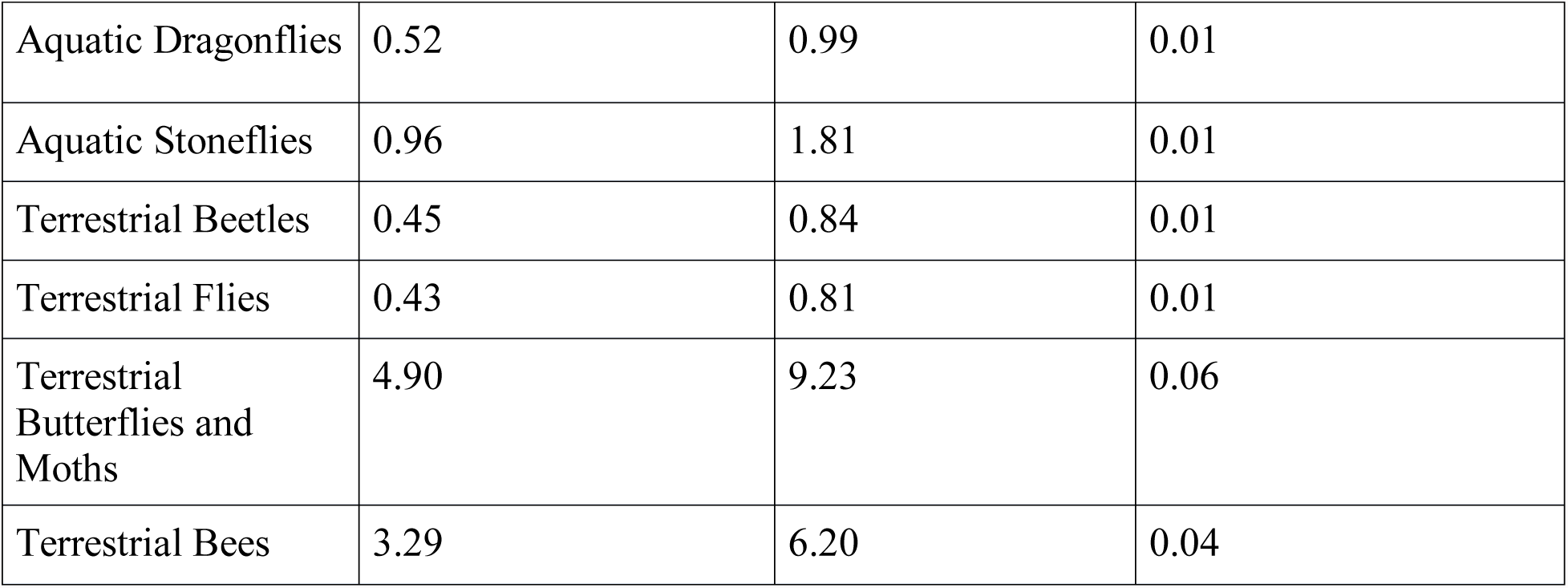
Potential EPA and DHA from ALA and potential total EPA at average, maximum, and minimum conversion efficiency. All data are expressed as percent of total fatty acids.

| **Food Item** | **EPA<sub>ALA</sub> Avg.** | **EPA<sub>ALA</sub> Max.** | **EPA<sub>ALA</sub> Min.** | **EPA<sub>Total</sub> Avg.** | **EPA<sub>Total</sub> Max.** | **EPA<sub>Total</sub> Min.** |
| --- | --- | --- | --- | --- | --- | --- |
| Aquatic Mayflies | 1.14 | 1.53 | 0.69 | 16.13 | 16.53 | 15.86 |
| Aquatic Dragonflies | 0.48 | 0.64 | 0.29 | 6.47 | 6.63 | 6.28 |
| Aquatic Stoneflies | 0.87 | 1.18 | 0.53 | 21.40 | 21.71 | 21.06 |
| Terrestrial Beetles | 0.41 | 0.55 | 0.25 | 5.18 | 5.32 | 5.02 |
| Terrestrial Flies | 0.39 | 0.53 | 0.24 | 2.83 | 2.97 | 2.68 |
| Terrestrial Butterflies and Moths | 4.47 | 6.01 | 2.69 | 6.65 | 8.19 | 4.87 |
| Terrestrial Bees | 3.00 | 4.03 | 1.81 | 3.00 | 4.03 | 1.81 |
| **Food Item** | **DHA<sub>ALA</sub> Avg.** |  | **DHA<sub>ALA</sub> Max.** |  | **DHA<sub>ALA</sub> Min.** |  |
| Aquatic Mayflies | 1.25 |  | 2.36 |  | 0.02 |  |

|  |  |  |  |
| --- | --- | --- | --- |
| Aquatic Dragonflies | 0.52 | 0.99 | 0.01 |
| Aquatic Stoneflies | 0.96 | 1.81 | 0.01 |
| Terrestrial Beetles | 0.45 | 0.84 | 0.01 |
| Terrestrial Flies | 0.43 | 0.81 | 0.01 |
| Terrestrial Butterflies and Moths | 4.90 | 9.23 | 0.06 |
| Terrestrial Bees | 3.29 | 6.20 | 0.04 |

**Table 3:**
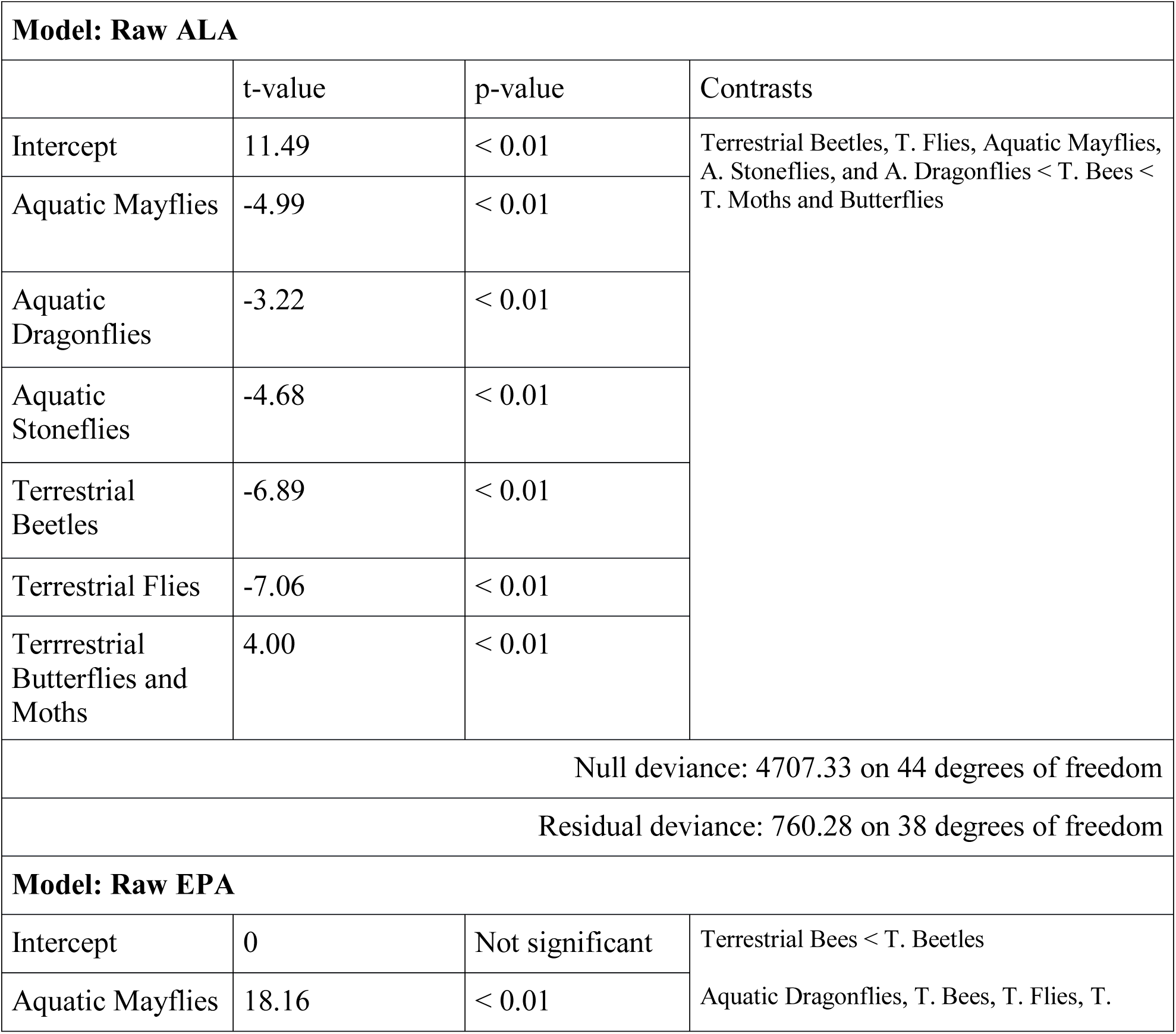
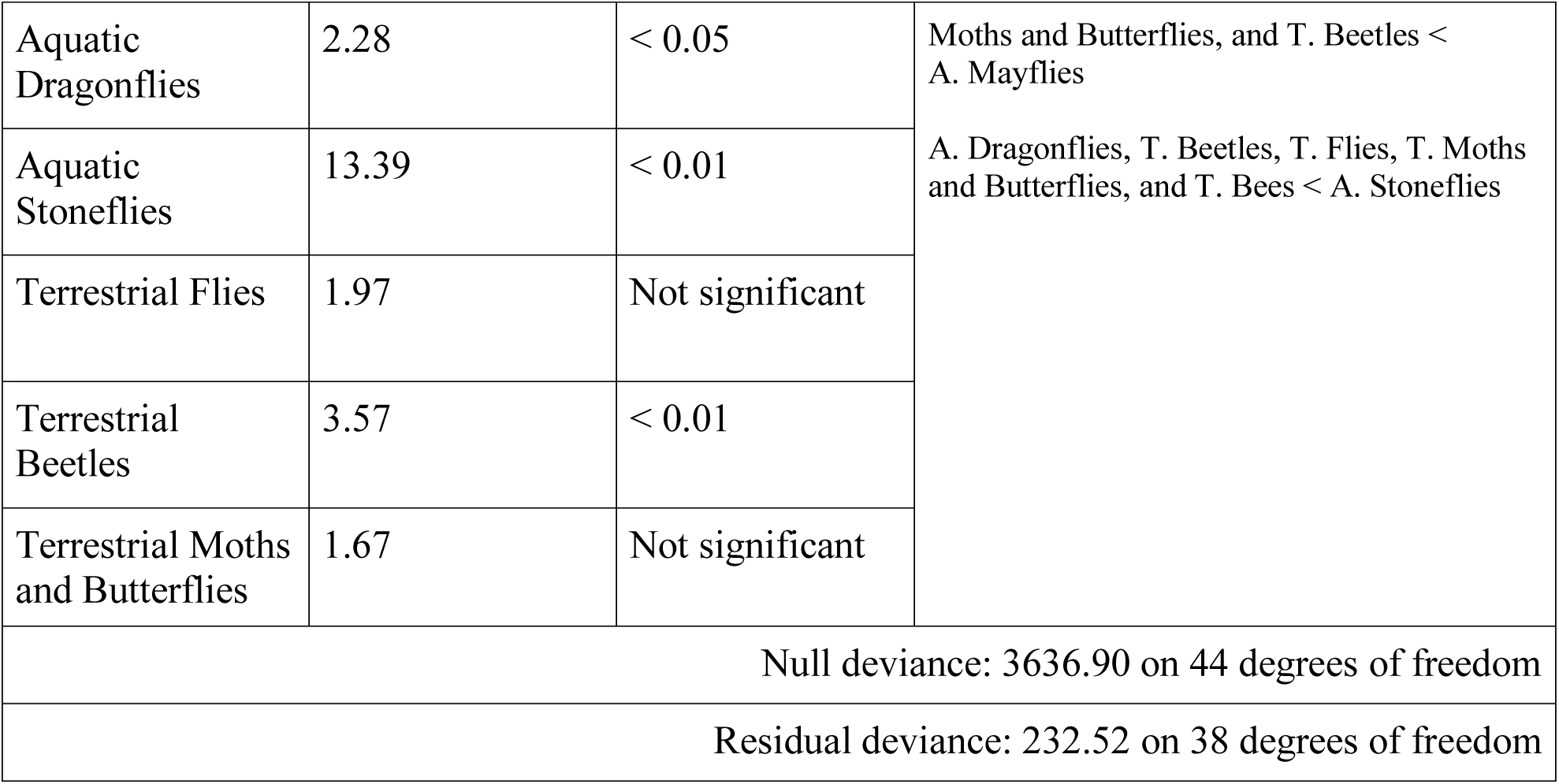
GLM results for raw ALA and EPA in insects

We estimated that Tree Swallows could derive significantly more potential EPA and DHA from ALA in terrestrial moths, butterflies, and bees than from other terrestrial or aquatic insects (Figure 1b; Table 4). However, total EPA (raw EPA plus potential EPA from ALA) from aquatic insects was still significantly higher than the largely ALA-derived total EPA from terrestrial moths, butterflies, and bees (two sample t-test for aquatic versus terrestrial total EPA: t-value = 8.05, df = 14.07, p-value < 0.01; Figure 1c; Table 5). Aquatic insect EPA derived primarily from diet (Figure 2) because aquatic insects had the significantly higher raw EPA values than did any terrestrial insects (Figures 1). Unlike aquatic insects, terrestrial insects, especially terrestrial moths, butterflies, and bees, had the potential to provide Tree Swallows with EPA primarily from ALA conversion (Figure 2).

**Figure 2:**
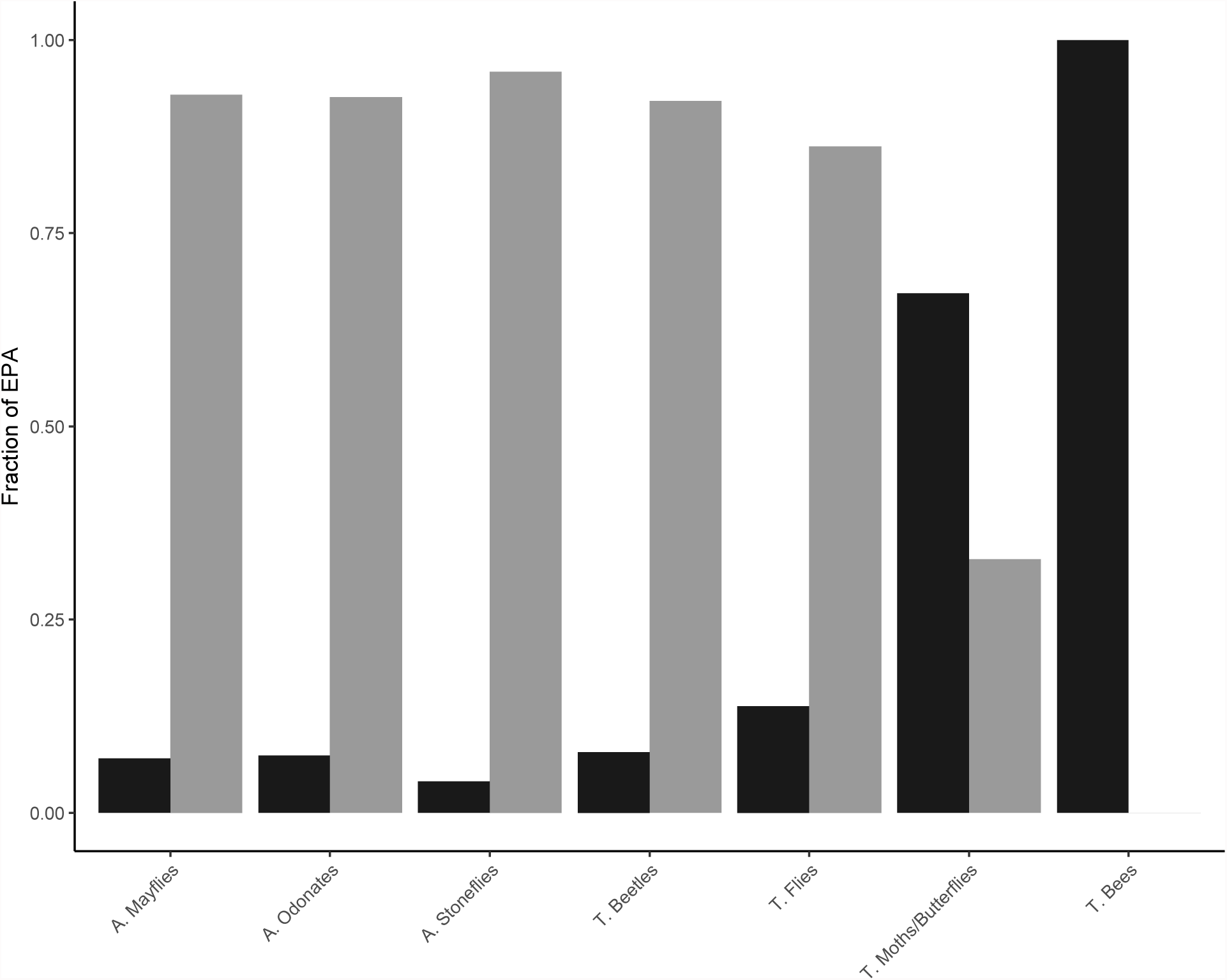
Fraction of EPA directly from diet or from ALA to LCPUFA conversion. LCPUFA derived from diet is gray and that from conversion is black.

**Table 4:**
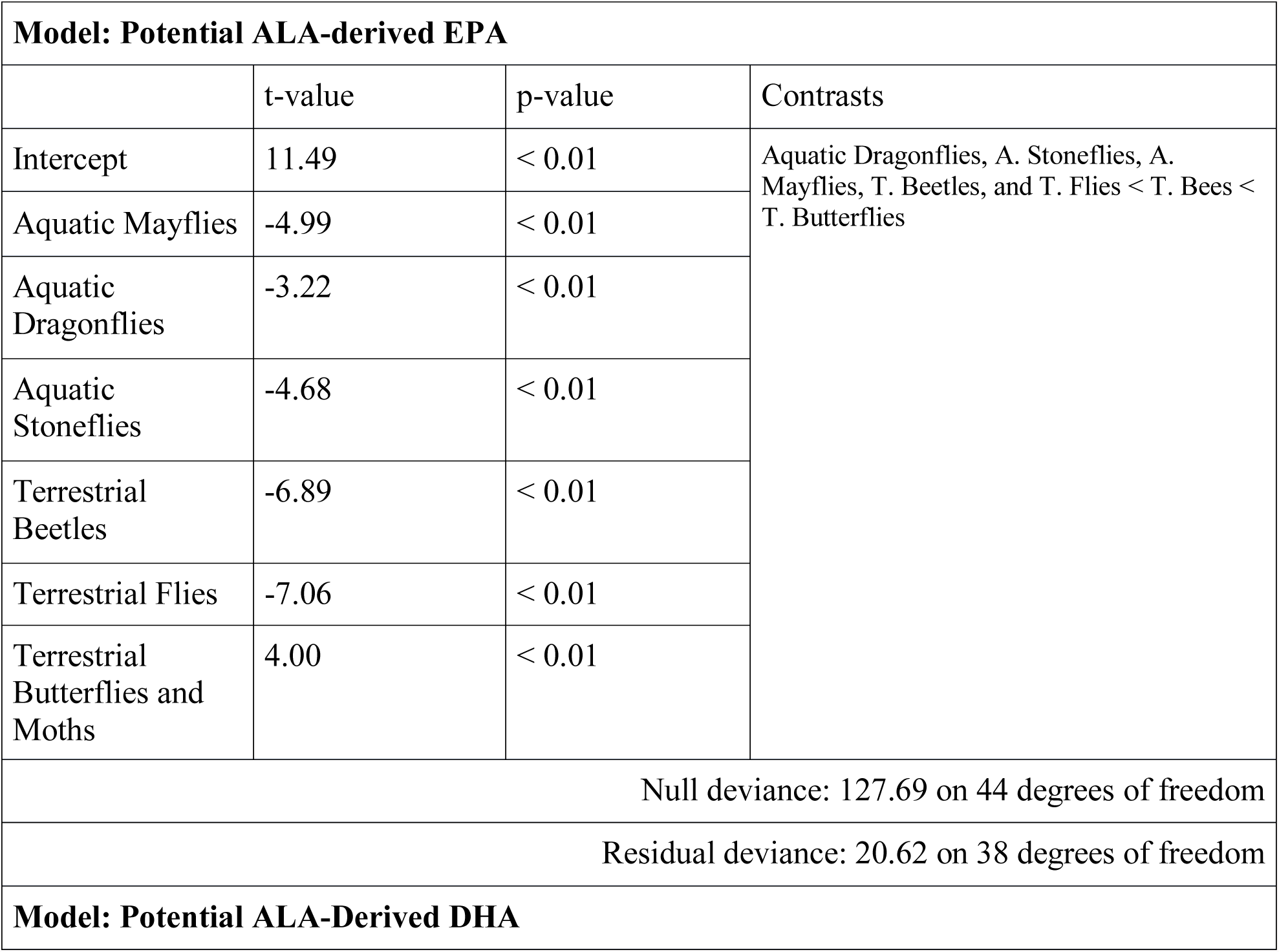
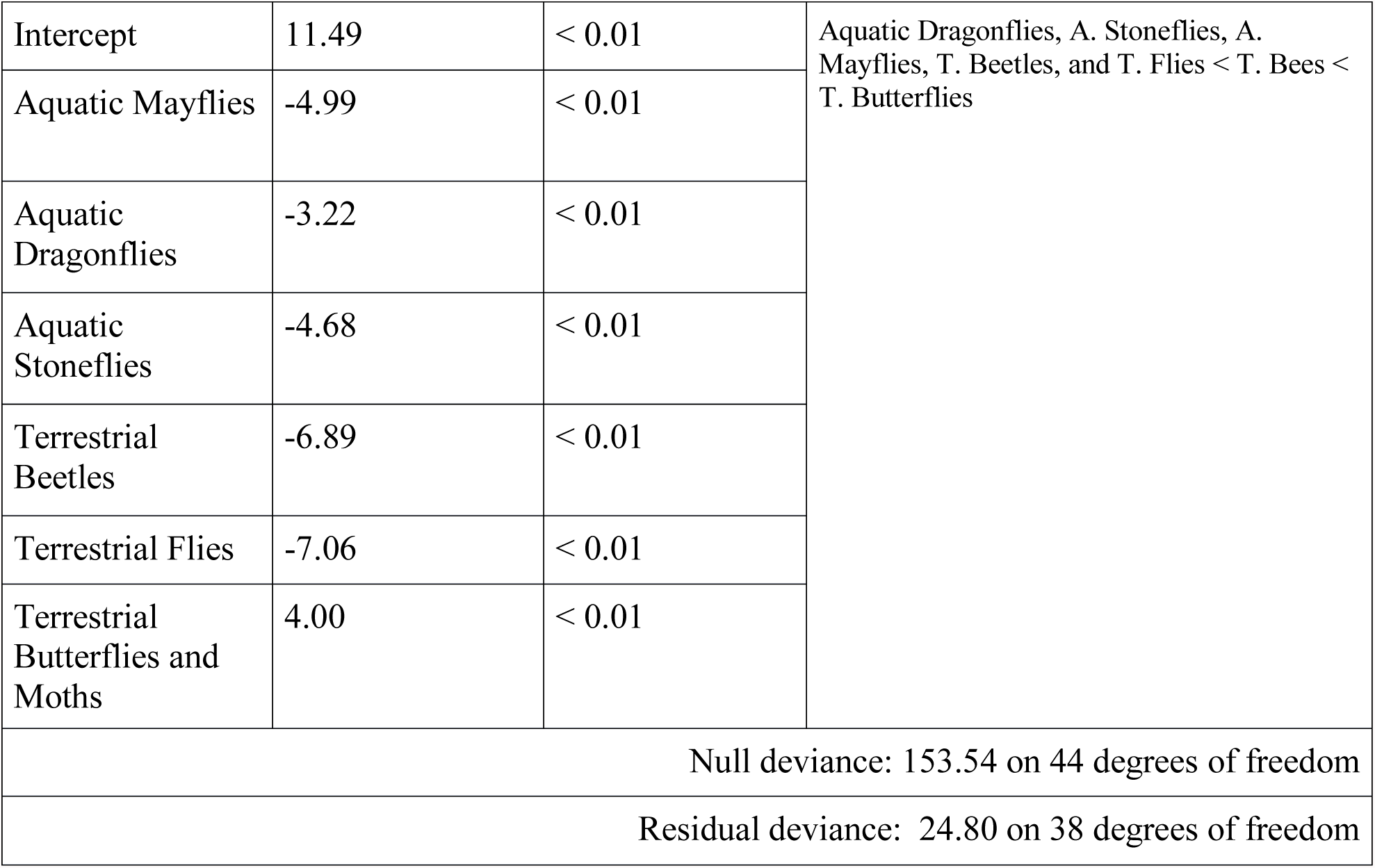
GLM results for potential ALA-derived EPA and DHA in insects

**Table 5:**
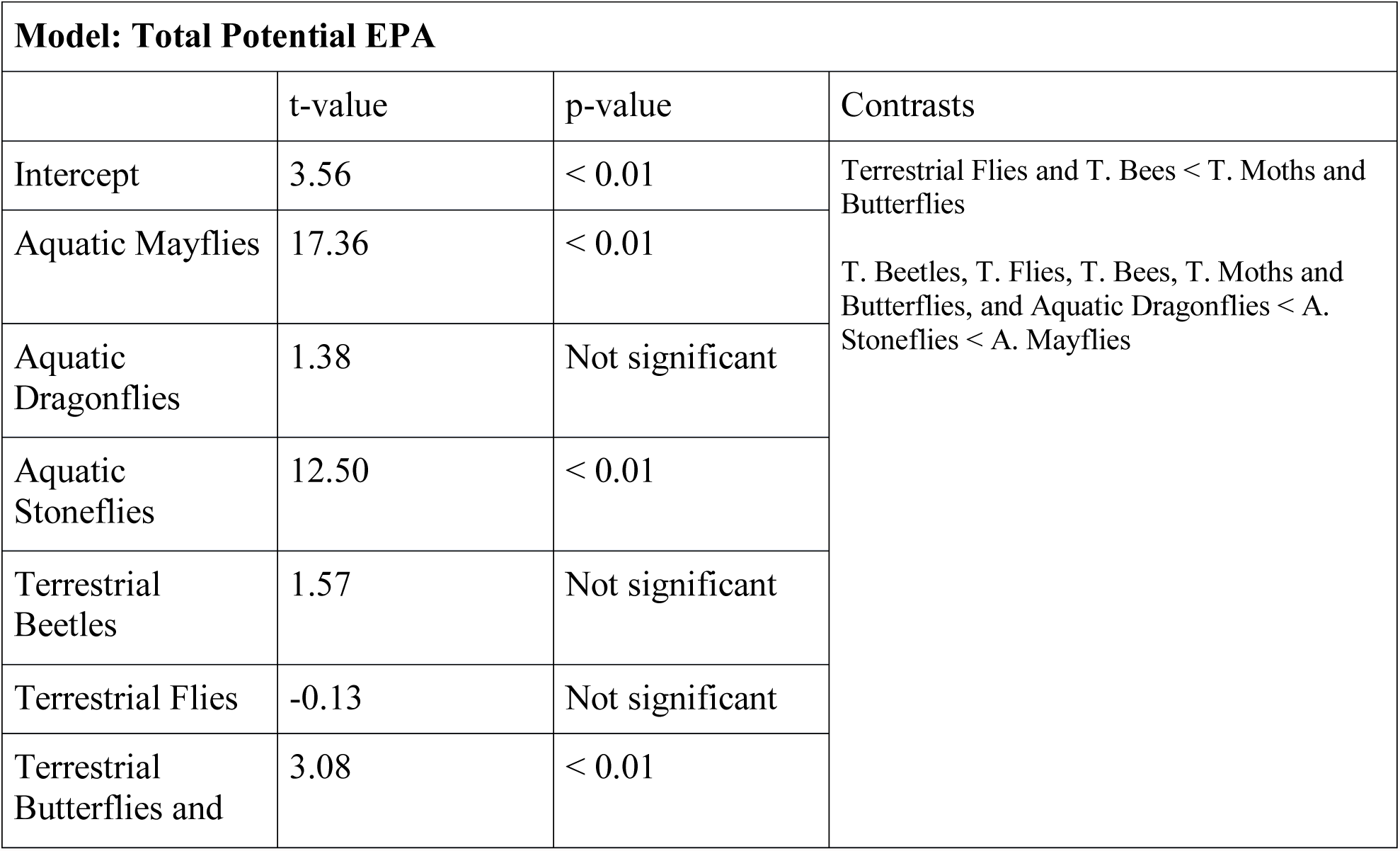
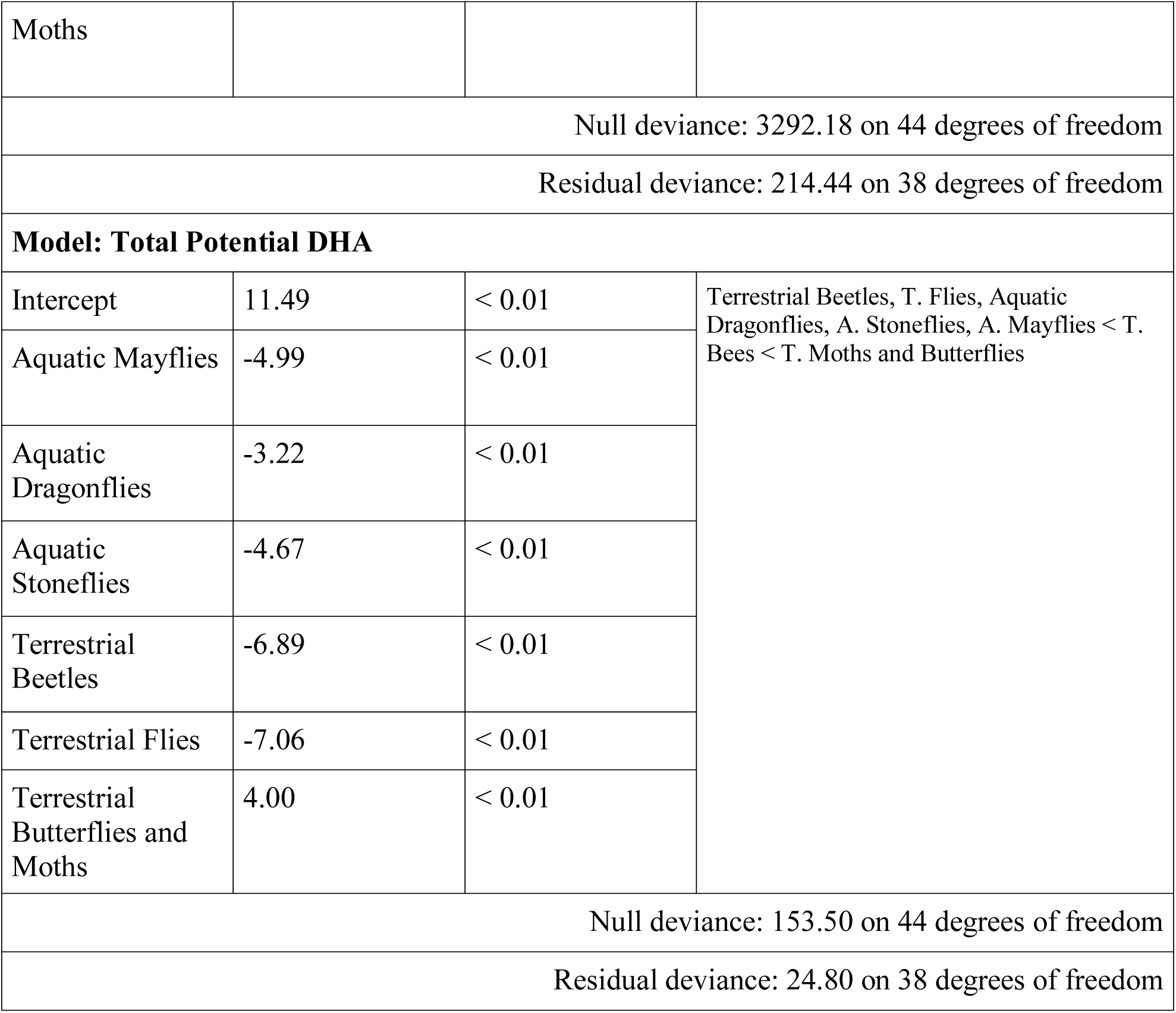
GLM results for potential total ALA and EPA in insects

In contrast to our findings for EPA, aquatic insects contained only trace amounts of DHA and no terrestrial insects contained any detectable DHA (Figure 1a). Therefore, regardless of conversion efficiency, over 95% of total DHA was derived from conversion, rather than diet for all insect prey (Tables 1-2). Due to their high ALA content, terrestrial moths, butterflies, and bees had the potential to supply significantly more DHA than aquatic insects or other terrestrial insects (Figure 1; Table 5). Our estimates for DHA from EPA-rich aquatic insects are likely conservative because our ^13^C-ALA tracer allowed us to measure ALA to DHA conversion as well as ALA-derived EPA conversion to DHA, but not direct EPA to DHA conversion (i.e., Tree Swallows could have converted additional unlabeled EPA to DHA).

## Discussion

Food quantity in terms of energy and food quality in terms of limiting nutrients are the major drivers of survival and performance in all living things everywhere, and especially in developing animals in the wild. We previously found that food containing LCPUFA reflective of aquatic insects improves multiple metrics of performance in Tree Swallow chicks in the laboratory (Twining et al. 2016a) and that the biomass of LCPUFA-rich aquatic insects is a strong predictor of Tree Swallow chick fledgling production in nature (Twining et al. submitted). In the present study, we sought to understand these costs to chick performance further by determining if and how Tree Swallow chicks can efficiently convert ALA into LCPUFA and deposit it into tissues. Our ALA tracer-based results reveal that Tree Swallow chicks are able to derive LCPUFA from ALA within liver and deposit ALA-derived LCPUFA into both liver and pectoral muscle. This evidence for ALA to LCPUFA conversion and deposition of ALA-derived LCPUFA suggests that performance costs in chicks on low LCPUFA diets (Twining et al. 2016a; Twining et al. in prep) likely arise from a combination of the energetic cost of converting ALA to LCPUFA as well as indirect LCPUFA limitation when ALA itself is in limited supply.

Our evidence that ALA to LCPUFA conversion is possible in Tree Swallows (Table 1) reveals that increased chick performance on diets with higher percentages of LCPUFA is likely due to a combination of direct LCPUFA limitation as well as the energetic savings accrued by receiving LCPUFA directly from of EPA-rich aquatic insects in diet. In our previous study, we found that Tree Swallow chicks suffered performance declines when feed on diets with 6.25% ALA, 1.47% EPA, and 1.42% DHA compared to diets with 1.82% ALA, 3.74% EPA, and 3.44% DHA (Twining et al. 2016a). While all insect taxa that we examined contain more than 1.82% ALA (the level of ALA in our higher performance dietary treatment), terrestrial Diptera provide less than the 3.74% EPA and 3.44% DHA levels in our high performance dietary treatment (Figure 1; Table 2), even at maximum measured deposition levels of 22.1% for EPA and 34.1% DHA (Table 2). Terrestrial insects that did contribute more total EPA than that of our high performing lab treatments provided substantially less EPA than aquatic insects (Figure 1). Only terrestrial butterflies and moths had the potential to provide more than sufficient total (raw plus potential from ALA) EPA and DHA (Figure 1). However, terrestrial butterflies and moths, beetles, and bees are all rare in Tree Swallow insect boluses fed to chicks, which are dominated by true flies (Diptera; McCarty and Winkler 1999; Winkler and others unpublished). Aquatic and terrestrial flies are likely the dominant prey in Tree Swallow boluses they are highly abundant and relatively easy to catch compared to less abundant and faster flying prey like butterflies (McCarty and Winkler 1999). Thus, it is highly unlikely that terrestrial insects alone can supply chicks with sufficient ALA-derive LCPUFA, making LCPUFA-rich aquatic insects ecologically essential for Tree Swallow chicks.

Strictly essential nutrients are nutrients that animals are unable to synthesize from their molecular precursors and must derive directly from diet. LCPUFA are thus not strictly essential for Tree Swallows because Tree Swallows can synthesize limited amounts of LCPUFA from precursors. However, LCPUFA-rich prey like aquatic insects appear to be crucially important, ecologically essential nutrients for Tree Swallows in natural systems because the ALA content of terrestrial insects and ALA to LCPUFA conversion efficiency are insufficient to supply chicks with the LCPUFA that they demand. In addition, ALA-derived LCPUFA levels were much higher in liver where much ALA to LCPUFA conversion occurs than in pectoral muscle (Table 1), where LCPUFA are deposited and used, but not synthesized. Our findings in Tree Swallows contrast with observations in chickens: chickens increase tissue LCPUFA with increased dietary LCPUFA, but are able to thrive even without dietary LCPUFA (Cherian and Sim 1991).

Past studies directly measuring ALA to LCPUFA conversion efficiency have focused on humans and other laboratory mammals (Brenna et al. 2009). Human studies uniformly show that ALA conversion to EPA is significant and in the few percent range, while conversion to DHA is barely above trace levels (Brenna et al. 2009). Our measured conversion efficiency in Tree Swallow liver (Table 1), where presumably most LCPUFA synthesis occurs, is broadly similar to that found in human blood (Burdge et al. 2004). However, ALA-derived LCPUFA levels in Tree Swallow pectoral muscle (Table 1), which is a more comparable tissue to blood in that LCPUFA synthesis is low or zero in both, were lower than levels in human blood (Burdge et al. 2004). Studies in rat pups suggest that ALA to LCPUFA conversion occurs at much higher efficiency than those we found in Tree Swallows even when LCPUFA are present in diet (Sheaff et al. 1995). However, although these studies suggest that ALA conversion is possible in humans and other mammals throughout their lifetime, most LCPUFA still come directly from dietary LCPUFA unless dietary sources are kept artificially low (Brenna et al. 2009).

Within the existing literature on ALA to LCPUFA conversion on humans, it is well known that ontogeny, sex, and reproductive status have important effects on conversion efficiency. For example, human females of reproductive age are more efficient at ALA to LCPUFA conversion than are human males of the same age (Burdge et al. 2002), which researchers have interpreted as an adaptation for provisioning human infants with LCPUFA (Brenna et al. 2009). Studies in humans also suggest that infants themselves are more efficient at conversion than are adults or older children, although infant conversion efficiency may be insufficient for optimal development in the absence of preformed dietary LCPUFA (Brenna et al. 2009). Even among human infants, those at earlier gestational ages appear to have higher LCPUFA conversion efficiency than do infants at later gestational ages (Carnielli et al. 2007).

Like human infants, the Tree Swallow nestlings that we studied are entirely reliant upon parental feeding. Nutritional demands are greatest for altricial temperate passerines like Tree Swallows during the nestling period when they undergo rapid growth, often doubling in mass every few days (Zach and Mayoh 1982). Thus, the conversion efficiency that we report for Tree Swallow nestlings is likely near maximum for the species under the diet conditions provided them. Unlike human mothers, who provision their fetuses and nursing infants with a constant supply of their own digested nutrients including LCPUFA (Brenna et al. 2009), or even precocial species of birds like chickens who provision embryos with high levels of ALA and LCPUFA in eggs (Lin et al. 1991; Speake and Wood 2005), wild altricial birds like Tree Swallows invest little LCPUFA into eggs (Speake and Wood 2005; Twining et al. in prep). Tree Swallows also complete nearly all somatic growth, including growing the brain, eyes, and other nervous tissues, within approximately three weeks. This means that nestling Tree Swallow chicks must acquire all of their LCPUFA within a very short time window, creating high selective pressure for high ALA to LCPUFA conversion efficiency and tissue deposition during the nestling period. As in humans, further studies on Tree Swallow adults are necessary to confirm how ALA to LCPUFA conversion efficiency varies throughout the life cycle. Tree Swallow chicks must be sexed with karyotypes and therefore we did not sex the chicks in this study. Studies comparing adult Tree Swallow males and females, which can be readily sexed in the field, will lend additional insights into the complete LCPUFA needs of these model avian insectivores throughout their life cycles.

Previous studies on birds and other animals in natural systems have not investigated ALA to LCPUFA conversion mechanistically using stable isotope tracers. Thus, unfortunately, it remains unclear if LCPUFA are either strictly or ecologically essential for many wild animals, including other birds. Most researchers have made inferences about conversion efficiency based on data on the effects of dietary LCPUFA content on performance or survival (e.g., Sargent et al. 1999) or tissue fatty acid composition (e.g., Cherian and Sim 1991). However, if conversion efficiency is low, costs to performance and survival may either be due to dietary LCPUFA limitation or the costs associated with converting ALA to LCPUFA. In contrast, if conversion efficiency is zero, then all costs to performance and survival must be due to direct limitation, making LCPUFA strictly essential. Direct measurements of conversion efficiency are clearly necessary to distinguish these possibilities.

Understanding whether LCPUFA are strictly essential, ecologically essential, or non-essential nutrients for birds and other wild animals is crucial for informing conservation efforts. In order to develop successful species management plans, environmental managers must understand the full suite of food and habitat resources that wild animals require throughout their life cycle. For example, our findings in Tree Swallow chicks suggest LCPUFA-rich aquatic insects and habitats are ecologically essential resources during a critical ontogenetic period. As a consequence, human activities that alter the composition and resulting nutritional quality of insect prey, such as land use change and pesticide use, as well as phenological shifts in insect emergence due to climate change have the potential to create nutritional mismatches for Tree Swallows (Twining et al. submitted). We hope that our field-based adaptation of an enriched stable isotope tracer method from human clinical studies is a starting point towards ultimately developing a general understanding of fatty acid nutritional ecology in wild animals.

## Acknowledgements

We thank Jeremy Ryan Shipley for field assistance and Tree Swallow nestling feeding advice. This work was approved under United States Fish and Wildlife Service migratory bird scientific collection permit #MB757670 and New York State Department of Environmental Conservation scientific collection permit #1477. All animal work was approved under Cornell Institutional Animal Care and Use Committee #2001-0051.

### Author Contributions

CWT and JTB conceived the ideas and designed methodology; CWT and PL collected the data; CWT, PL, and JTB analyzed data; CWT led the writing of the manuscript and JTB, ASF, and DWW contributed critically to the drafts and all authors gave final approval for publications.

### Competing Interests

No competing interests declared.

### Funding

CWT, PL, and JTB acknowledge funding from USDA Hatch Grant NYC-399-7461; CWT and ASF acknowledge funding from the NSF DDIG DEB-1838331; and CWT acknowledges funding from a Cornell University College of Agriculture and Life Sciences Mellon Grant. CWT was supported by the NSF GRFP during the course of this project.

